# Honey bee pollen foraging ecology across an urbanization gradient

**DOI:** 10.1101/824474

**Authors:** Rodney T. Richardson, Tyler D. Eaton, Chia-Hua Lin, Garrett Cherry, Reed M. Johnson, Douglas B. Sponsler

## Abstract

Understanding animal foraging ecology requires large samples sizes spanning broad environmental and temporal gradients. For pollinators, this has been hampered by the laborious nature of morphologically identifying pollen. Metagenetic pollen analysis is a solution to this issue, but the field has struggled with poor quantitative performance. Building upon prior laboratory and bioinformatic methods, we applied quantitative multi-locus metabarcoding to characterize the foraging ecology of honey bee colonies situated along an urban-agricultural gradient in central Ohio, USA. In cross-validating a subset of our metabarcoding results using microscopic palynology, we find strong concordance between the molecular and microscopic methods. Our results show that, relative to the agricultural environment, urban and suburban environments were associated with higher taxonomic diversity and temporal turnover of honey bee pollen forage. This is likely reflective of the fine-grain heterogeneity and high beta diversity of urban floral landscapes at the scale of honey bee foraging. Our work also demonstrates the power of honey bees as environmental samplers of floral community composition at large spatial scales, aiding in the distinction of taxa characteristically associated with urban or agricultural land use from those distributed ubiquitously across our landscape gradient.

## Introduction

As a perennial, eusocial and cosmopolitan florivore, the western honey bee (*Apis mellifera* L.) faces unique demands for resource acquisition. These demands are met by a sophisticated colony-level foraging biology characterized by extreme dietary generalism, energetic optimization, and the integration of private and shared information about landscape-scale floral resources (Seeley 1995, Grüter and Farina 2009). These traits make honey bee foraging responsive to daily and seasonal turnover of floral resources within a given landscape (Visscher and Seeley 1982, Couvillon et al. 2014b) and adaptable to the floristic variation across landscapes ranging from deserts to rainforests in every continent but Antarctica. Nevertheless, honey bee fitness responds to landscape composition (Lecocq et al. 2015, Sponsler and Johnson 2015, Smart et al. 2016, Döke et al. 2018), indicating that behavioral sophistication cannot fully compensate for variation in landscape-scale floral resource availability. This inference is corroborated by empirical and theoretical studies demonstrating that the adaptive value of the waggle dance—the pinnacle of honey bee foraging sophistication—varies with the spatiotemporal distribution of floral resources (Donaldson-Matasci and Dornhaus 2012, Schürch and Grüter 2014).

During the Anthropocene, honey bee biogeography has come to include the novel landscapes created by human agriculture and urbanization, both of which generate floristic conditions that are far-removed from their nearest natural analogs. In typical agricultural landscapes, large crop monocultures are the predominant feature, intercalated with marginal non-crop habitat features like field margins, roadsides, and riparian buffers. In contrast, urban landscapes are characterized by an extensive impervious matrix finely intermixed with relatively small patches of greenspace composed of a mixture of native and exotic species, in some cases resulting in a net species richness equal to or greater than that of nearby non-urban areas, though typically with lower species density (McKinney 2008, Aronson et al. 2014, Aronson et al. 2015). Ultimately, agricultural and urban landscapes diverge substantially both from each other and from the historic landscapes they displace.

The question of how honey bee foraging behavior responds to the unique challenges of urban and agricultural landscapes is salient from both basic and applied perspectives (Couvillon et al. 2014a, Härtel and Steffan-Dewenter 2014). With respect to urban land use, recent empirical studies have described honey bee (Lau et al. 2019, Lucek et al. 2019) diet and foraging distance (Garbuzov et al. 2015). The only study, however, to articulate a theoretical relationship between honey bee foraging behavior and suburban land use is that of Waddington et al. (1994), where the authors suggest that the fine-grain pattern of floral patch distribution typical of suburban landscapes favors independent patch discovery over the use of waggle dance recruitment by foraging honey bees. Under this hypothesis, researchers would expect a greater number of floral patches to be simultaneously exploited in suburban areas. With respect to agricultural landscapes, Steffan-Dewenter and Kuhn (2003) suggest that highly simplified agricultural landscapes with large monoculture patches result in fewer patches being simultaneously exploited, and Nürnberger et al. (2019) conclude that patch discovery would be more dependent on waggle dance recruitment in this circumstance. Together, these studies formulate essentially reciprocal hypotheses regarding the effects of landscape on optimal foraging strategy. Thus, a preliminary synthesis of existing literature would be the hypothesis that urban and agricultural land use have divergent effects on honey bee spatial foraging patterns, the former generating more diffuse foraging and the latter generating a more clumped recruitment-driven foraging. A logical extension of this spatial hypothesis would be the prediction that the taxonomic composition of honey bee foraging would follow similar patterns, being more diversified in urban landscapes and more homogenized in agricultural landscapes.

Areas where agricultural and urban landscapes occur in close proximity present ideal opportunities to study the behavioral response of honey bee foraging to the respective challenges presented by these landscapes while controlling for regional climate and floristics (Sponsler et al. 2017). In the present study, we employ DNA metabarcoding to identify the botanical origins of honey bee-collected pollen sampled along an urban-agricultural gradient in central Ohio, USA. By characterizing the taxonomic composition, diversity, and turnover of these pollen samples, we revisit the hypotheses of Waddington et al. (1994) and Steffan-Dewenter and Kuhn (2003), testing the prediction that, along a gradient spanning from fine-grain urban land use to highly simplified agricultural land use, pollen diversity and sample-to-sample turnover will correlate positively with increasing urbanization.

## Methods

### Site selection and sample collection

In the spring of 2013, we established apiaries at four sites spanning an urban-agricultural gradient (Figure 1). Two of these sites (BK and GH) were within the metropolitan area of Columbus Ohio, USA, with a third apiary (CC) residing on the edge of a western suburb and the fourth apiary (FSR) residing approximately 23 km from the western edge of the city. Each apiary contained two to six eight-frame Langstroth hives, two of which were fitted with Sundance I bottom□mounted pollen traps (Ross Rounds, Albany, NY, USA). From late spring to late fall of both 2013 and 2014, we continuously trapped corbicular pollen from one hive per apiary on a weekly basis, collecting 109 bulk pollen samples in total. To avoid causing pollen starvation, we alternated trapping between colonies from one sampling event to the next.

**Figure 1:**
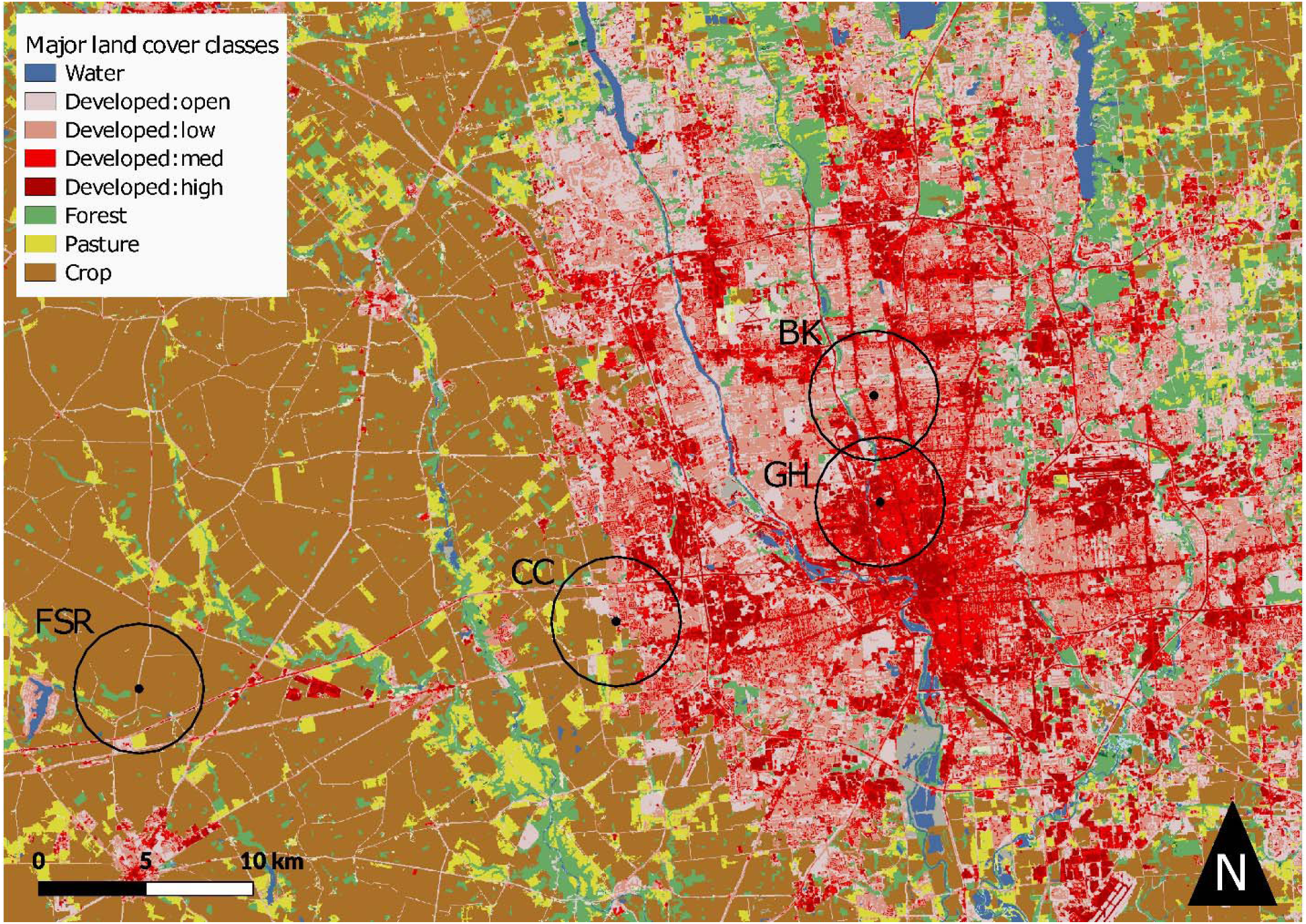
A landcover map of Columbus, OH and surroundings. Black dots indicate apiary locations and circles are drawn at a 3 km radius surrounding each apiary.

### GIS landscape analysis

We analyzed landscape composition using the 2011 National Land Cover Database (NLCD) land cover layer (Homer et al. 2015). For each site, we extracted from the NLCD layer a 3 km radius and tabulated its composition using QGIS (Team 2015) and R (Team 2014). As an approximate measure of the degree of urbanization for each site, we summed all four categories of developed land classifications of the NLCD layer into a single ‘urban’ category. This measure of urbanization ranged from 7 to 98 percent of the total land area within the landscapes surrounding our sites.

### Microscopic palynology

For a subset of our samples, 9 samples collected at site CC between July 31st and September 26th of 2014, microscopic palynology was used to characterize pollen taxonomic composition. These microscopy results were published in a previous study (Sponsler et al. 2017), otherwise independent from the present work. Briefly, bulk pollen samples were subsampled to 10 percent of their total weight up to a maximum of 10 g for samples with mass of greater than 100 g. Subsamples were then sorted by color and texture, mounted with fuchsin-stained glycerin jelly, and identified by comparison to a local reference collection. Proportional abundance of taxa within each sample were calculated using the volumetric approach of O’Rourke and Buchmann (1991).

### Molecular pollen analysis laboratory procedures

Broadly, our molecular pollen analysis methods follow those of Richardson et al. (2019). Prior to DNA extraction, corbicular pollen pellets were subsampled and homogenized using previously described methods. For each sample, the lesser amount of either 20 g or 10 percent by mass of each bulk sample was added to distilled water to yield a final concentration of 0.1 g/mL of pollen. The pollen mixture was placed in a blender (Hamilton Beach #54225, Southern Pines, NC, USA) and homogenized for 2.5 min. Pollen homogenate, 1.4 mL, was then placed in a 2 mL bead beater microfuge tube (Fisherbrand Free□Standing Microcentrifuge Tubes; Fisher Scientific, Hampton, NH, USA). To facilitate mechanical pollen disruption during bead beating, 3,355 mg of 0.7 mm zirconia beads (Fisher Scientific, Hampton, NH, USA) were added to each tube. Tubes were placed in a bead-beater (Mini□BeadBeater□ 16; BioSpec Products, Bartlesville, OK, USA) for 5 minutes at the highest setting. Samples were briefly agitated with a vortex mixer prior to removing 400 µl of homogenate for DNA extraction using the DNeasy Plant Minikit (QIAGEN Inc., Valencia, CA) according to the manufacturer’s protocol.

Three loci were selected for amplicon library preparation: *rbcL, trnL* and ITS2. For compatibility with the Illumina Nextera sequencing protocol, we conducted a 3-step PCR protocol in which the product of each step was used as the template in the next step. Since this protocol was necessary for each locus, each sample required a total of nine PCR reactions. The following templates were used sequentially in the three PCR steps: 1) 100–150 ng of pollen DNA extract; 2) 1 µl of unpurified PCR product from the first PCR reaction; 3) 1 µl of unpurified PCR product from the second PCR reaction. Universal primers (White 1990, Taberlet et al. 1991, Kress and Erickson 2007, Palmieri et al. 2009, Chen et al. 2010) with no 5-prime fusion oligos were used in the first PCR reaction. The primers used in the second and third PCR reactions were modified to allow template priming, sample indexing, and lane hybridization during Illumina sequencing. Each of the three PCR steps were performed in a standard 20 µl reaction containing 10 µl of Phusion Flash High-Fidelity PCR Master Mix (Thermo Fisher Scientific, Inc), template DNA, and a pair of DNA primers. After the third PCR, amplicon products were purified and normalized using the SequalPrep Normalization Plate Kit (Thermo Fisher Scientific, Waltham, MA, USA), pooled equimolarly and sequenced using Illumina MiSeq (2 × 300 cycles). Further details on primer sequences, PCR conditions, and sample dual-indexing design are provided at https://github.com/RTRichar/UrbanRuralPollen.

### Taxonomic annotation of sequence data

During bioinformatic processing, mate-paired sequences were first merged using Pear (v.0.9.1; (Zhang et al. 2013)). While *rbcL* and ITS2 were short enough for mate-pair merging, the *trnL* amplicon we used often exceeded 600 bp in length and few sequences could be merged for this locus. Thus, we used the Metaxa2 Database Builder (v1.0 beta 4; (Bengtsson-Palme et al. 2018)) along with the Metaxa2 classifier (v2.2; (Bengtsson-Palme et al. 2015)) to identify *trnL* sequences, reverse-complement the reverse reads and concatenate the mate-pairs into contiguous sequences. Semi-global top-hit alignment was then performed with VSEARCH (v2.8.1; (Rognes et al. 2016)) to assign taxonomy to all merged or concatenated sequences using custom reference sequence databases, geographically filtered to only include sequences belonging to genera known to be present in Ohio according to the USDA Plants Database. These databases were curated using MetaCurator (v1.0.1; (Richardson et al. 2019)), which utilizes HMMER (v3.1; (Eddy 2011)), MAFFT (v7.270 (Katoh et al. 2002)) and VSEARCH to extract reference sequences of the amplicon barcode of interest from all available sequences, including whole genome assemblies.

### Statistical analysis of sequencing results

Upon taxonomically annotating our sequence data, we employed the consensus-filtered, median-based analysis implemented in Richardson et al. (2019). Briefly, for each sample, we excluded genera identified with only one of the three barcode loci. We then calculated the median proportional abundance of these consensus-filtered genera while excluding zeros as non-detections. We then regressed the results of each individual locus, as well as the metabarcoding median against the microscopic data using least squares regression to characterize their quantitative performance. For these regressions, genus-level proportions were used for all genera except those from the family Asteraceae, which were summed to the family level due to the difficulty of morphologically distinguishing these genera.

In addition to testing the molecular data against the microscopic data for a subset of our samples, we investigated the relationship between ITS2-derived taxonomic proportions and metabarcoding median proportions at the genus level for all samples. Given previous studies (Keller et al. 2015, Richardson et al. 2015, Smart et al. 2017, Bell et al. 2019), it was surprising that ITS2 showed the best agreement with the microscopic validation data, and we sought to test if the experiment-wide ITS2 proportions were in strong disagreement with the three-marker median estimates.

We chose to conduct ecological inferences using 3,000 randomly sampled ITS2 sequences per sample as opposed to using the taxonomic proportions generated using the consensus-filtered metabarcoding median results for the following reasons. First, the ecological indices estimated were dependent upon normalized sampling effort, an assumption which was not feasible across all three markers since several *trnL* libraries failed to yield sufficient coverage for analysis, despite the fact that we analyzed all libraries with gel electrophoresis to ensure the presence of the appropriately sized product prior to sequencing. Such library dropout is not uncommon among plant metabarcoding literature (Sickel et al. 2015), and likely points to amplicon clustering issues or an insufficient number of PCR amplification cycles for this particular marker. Second, genus-level ITS2 taxonomic proportions were strongly correlated with both the microscopic palynology results as well as the three-marker median proportions, indicating that they provided sufficient characterization of our samples both quantitatively and qualitatively.

Using the standardized ITS2 dataset, we estimated genus-level Shannon diversity using the Vegan R package (Dixon 2003). We also estimated two indices of temporal turnover in forage at the genus level, mean rank shift (MRS), a non-parametric index of the degree of quantitative change from one sampling to the next, and qualitative turnover, the sum of taxon appearances and disappearances between sampling events. Qualitative turnover was calculated with a custom R script, while MRS was estimated using the Codyn R package (Hallett et al. 2016). The commands, R and python code, and reference sequence databases used for bioinformatic analysis can be found at https://github.com/RTRichar/UrbanRuralPollen.

With ITS2-based estimates of sample diversity, MRS and qualitative turnover, we calculated the median of each ecological index for each site and year. We then regressed these annual index medians against the percent developed land within the surrounding landscape using ordinary least squares regression. For these regressions, we calculated one-tailed *p*-values as we expected higher diversity and temporal turnover at the urban extreme of the gradient at the outset of our experiment. Further, we conducted non-metric multidimensional scaling (NMDS) of these standardized ITS2 datasets to visualize differences in forage composition by site and year. For this analysis, the ggrepel R package (Kamil Slowikowski 2019) was used to dodge text elements.

## Results

### Sequencing run statistics

After sequencing, we obtained 4,024,124 mate-paired reads, of which 3,094,447 were clearly identifiable as belonging to Viridiplantae after VSEARCH alignment. On average, individual amplicon libraries were sequenced to a depth of 9,463 ± 286 SEM, although there was significant variation in the number of sequences per library across the three markers (*P* < 0.001, one-way ANOVA), with the mean and standard error of sequences obtained per library for *rbcL, trnL* and ITS2 being 9,862 ± 422, 7,439 ± 626 and 11,088 ± 340, respectively. A follow-up Tukey’s HSD test revealed a significant decrease in sequencing depth for *trnL* libraries in particular (*P* < 0.005 for both the *trnL*-*rbcL* and *trnL*-ITS2 comparisons).

### Quantitative evaluation of pollen metabarcoding data

Regressing the cube-root squared metabarcoding proportions against the cube-root transformed microscopic palynology proportions (Figure 2), we found strong and significant agreement between the two methods for ITS2 (*P* < 0.001, *R*^2^ = 0.72) and the three-marker metabarcoding median (*P* < 0.001, *R*^2^ = 0.61). Further, significant relationships were found for *trnL* and *rbcL*, though these relationships were considerably weaker (*P* < 0.001 for both markers, *R*^2^ = 0.41 for *rbcL, R*^2^ = 0.21 for *trnL*). Notably, while the untransformed molecular and microscopic proportions were also highly correlated (*P* < 0.001 for all four tests), the differential transformations we report above resulted in considerably improved model fit, with 4 to 16 percent more variance explained across the four model comparisons, indicating that the molecular results exhibited a slight non-linear bias. Lastly, analysis of the relationship between ITS2 and the three-marker median proportions revealed a strong correlation (*P* < 0.001, *R*^2^ = 0.71).

**Figure 2:**
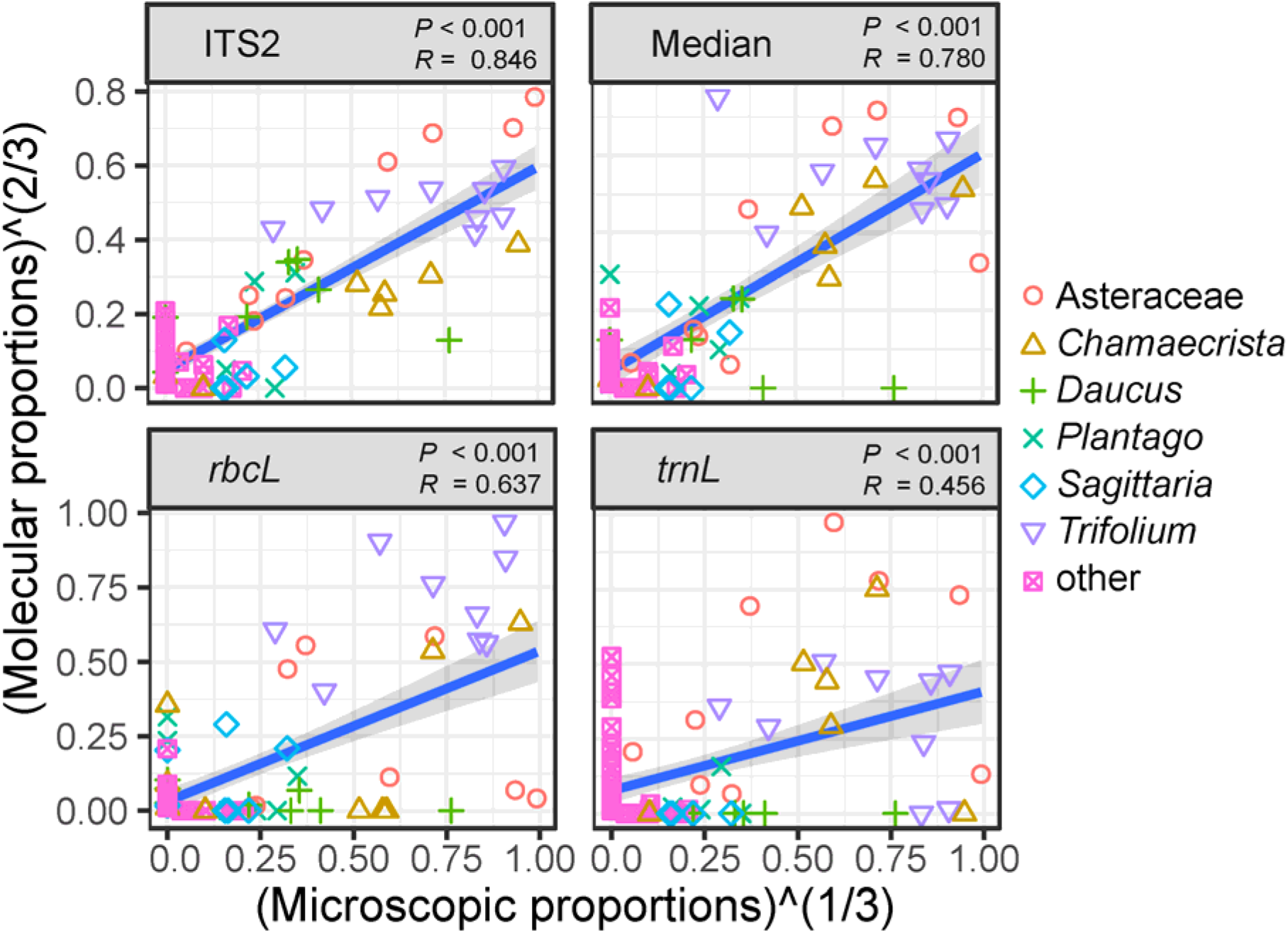
Molecular proportions regressed against microscopic proportions for the nine samples analyzed by pollen morphology. Taxa are shown with unique colors and point types with the exception of minor taxa, those which never exceeded 2.5 percent relative abundance according to microscopy. The comparison was performed at the genus level with the exception of Asteraceae, for which genera are difficult to distinguish morphologically.

### Pollen forage composition across the urban-agricultural gradient

Our results indicate that plant species within Fabaceae and Asteraceae, predominantly *Trifolium, Chamaecrista, Solidago, Symphyotrichum* and *Eurybia*, provision the majority of the summer and fall honey bee pollen diet at our study sites (Figure 3 and Figure 4). On average, these two families accounted for a mean and standard error of 73 ± 2 percent across all samples analyzed, with the genus *Trifolium* comprising an average and standard error of 36 ± 2 percent of each sample.

**Figure 3:**
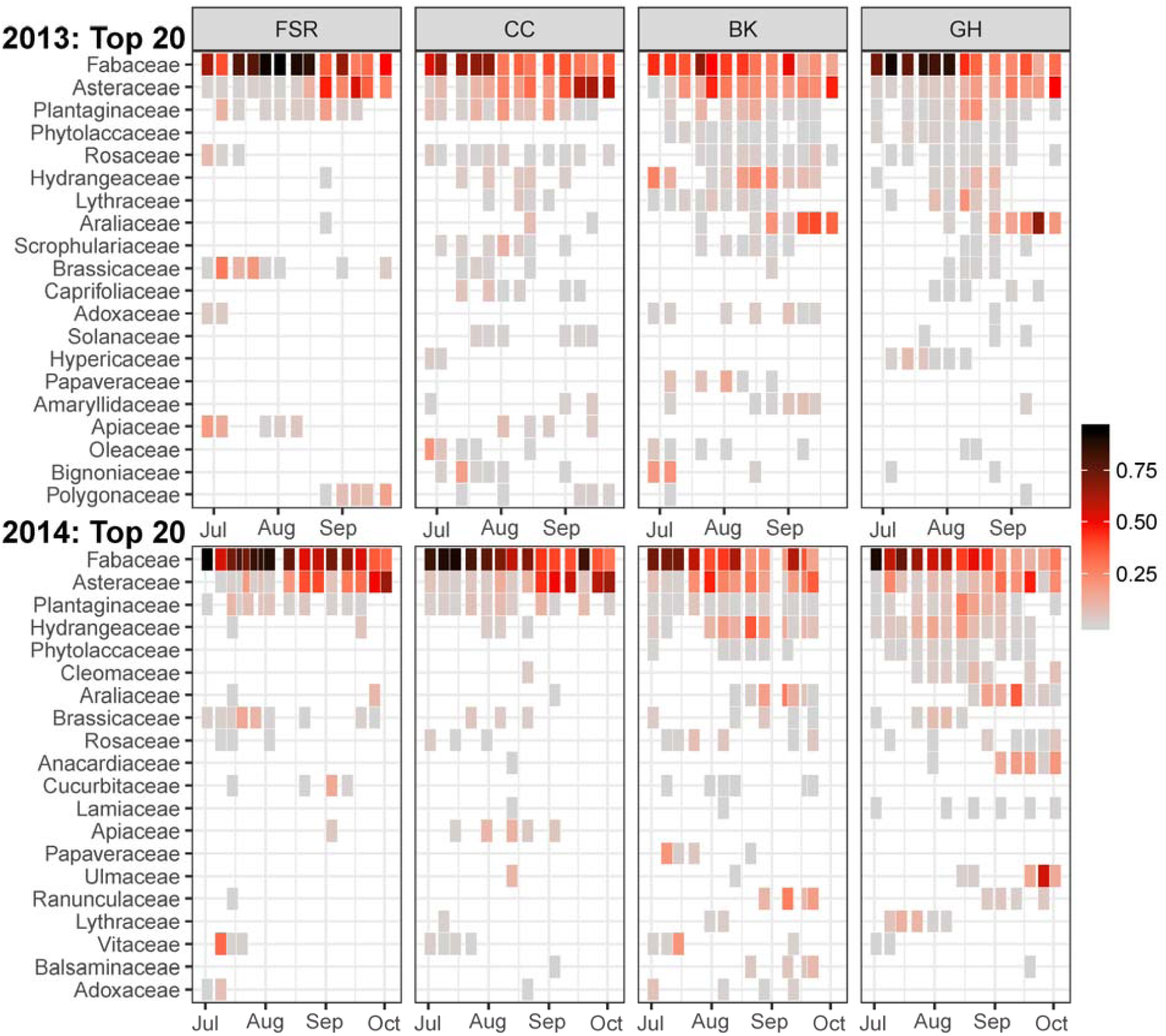
Proportional abundances of top 20 families detected, as measured by mean frequency of occurrence across sites for 2013 and 2014 separately. Families are ordered by decreasing frequency of detection for visualization purposes. Panels are arranged from the most agricultural site (FSR) to the most urban site (GH).

**Figure 4:**
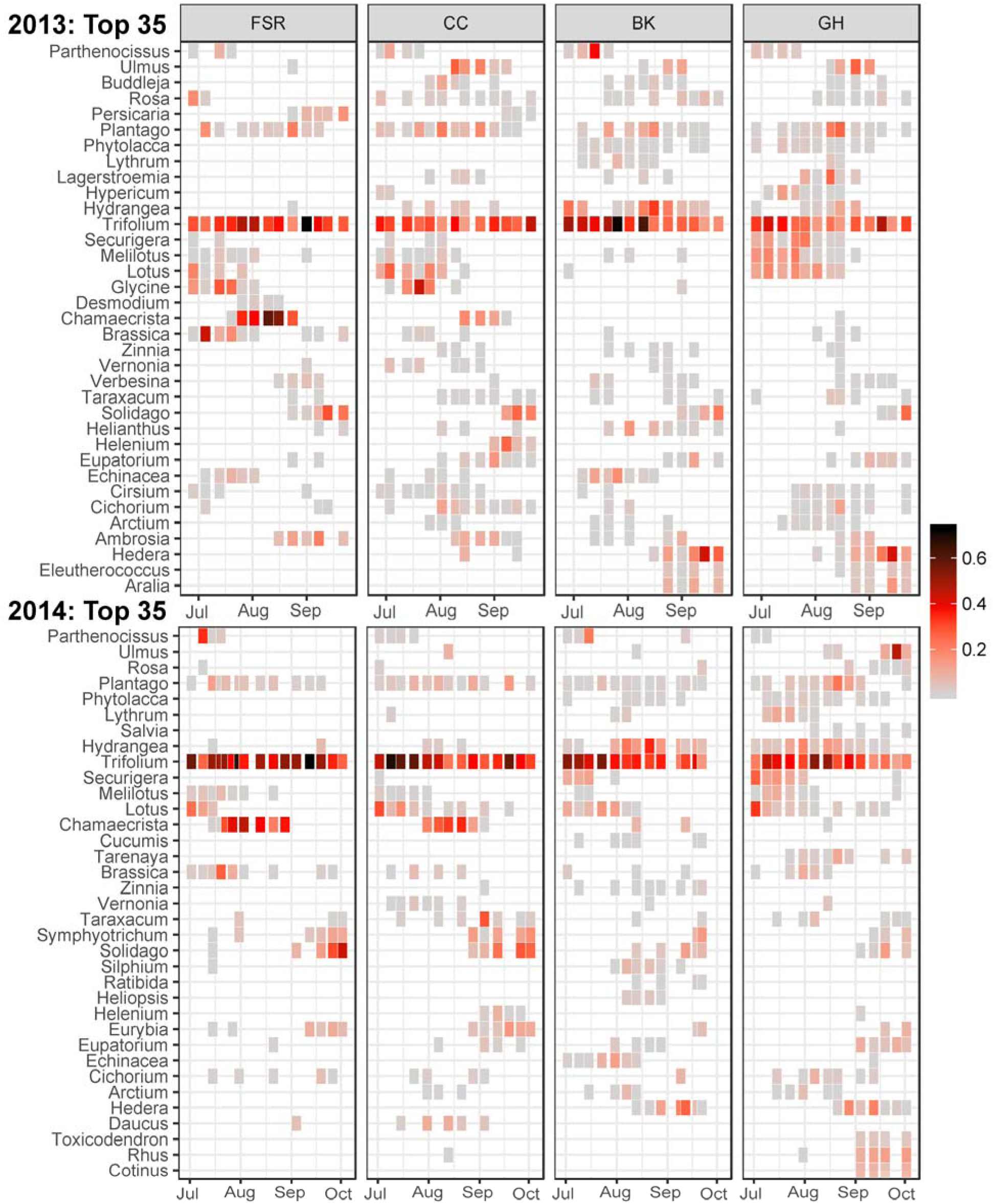
Proportional abundances of the top 35 genera detected, as measured by mean frequency of occurrence across sites for 2013 and 2014 separately. Genera are grouped by family for visualization purposes. Panels are arranged from the most agricultural site (FSR) to the most urban site (GH).

Across the landscape gradient, we detected honey bee foraging on taxa characteristically associated with either urban or agricultural landscapes (Figure 4). Apiaries with access to rural agricultural land cover collected small proportions of *Zea mays* pollen and large proportions of *Glycine, Brassica* and *Chamaecrista*. Urban and suburban apiaries appeared to be associated with common ornamental and urban-naturalized plants such as *Ulmus, Hedera, Hydrangea* and *Buddleja*. We also observed pollen foraging on disturbance-associated taxa, such as *Aralia, Rhus, Arctium* and *Phytolacca*, toward the urbanized end of the gradient.

### Ecological inference across the urban-agricultural gradient

Analyzing median estimates of diversity, mean rank shift (MRS), and qualitative turnover per site and per year, revealed interesting trends (Figure 5). For 2013, diversity, MRS and qualitative turnover were apparently higher in the urban relative to the agricultural site, with the highest median of each index observed in the intermediate half-suburban and half-agricultural site, CC. However, none of these ecological indices were significantly linearly related to the proportion of developed land within the landscape surrounding each apiary (*P* > 0.1 for all tests). In 2014, we observed the same trend, wherein higher ecological index median values were observed toward the urban end of the gradient. For 2014, each of the three indices were significantly linearly related to the proportion of developed land surrounding each apiary (*P* = 0.027 for diversity, *P* = 0.004 for MRS and *P* = 0.005 for qualitative turnover). Additionally, the two-year average of median diversity per site was significantly related to the proportion of developed land surrounding each apiary (*P* = 0.003). The two-year averages of median MRS and qualitative turnover were not significantly linearly related to the proportion of developed land surrounding each apiary (*P* = 0.097 and *P* = 0.123 for MRS and qualitative turnover, respectively).

**Figure 5:**
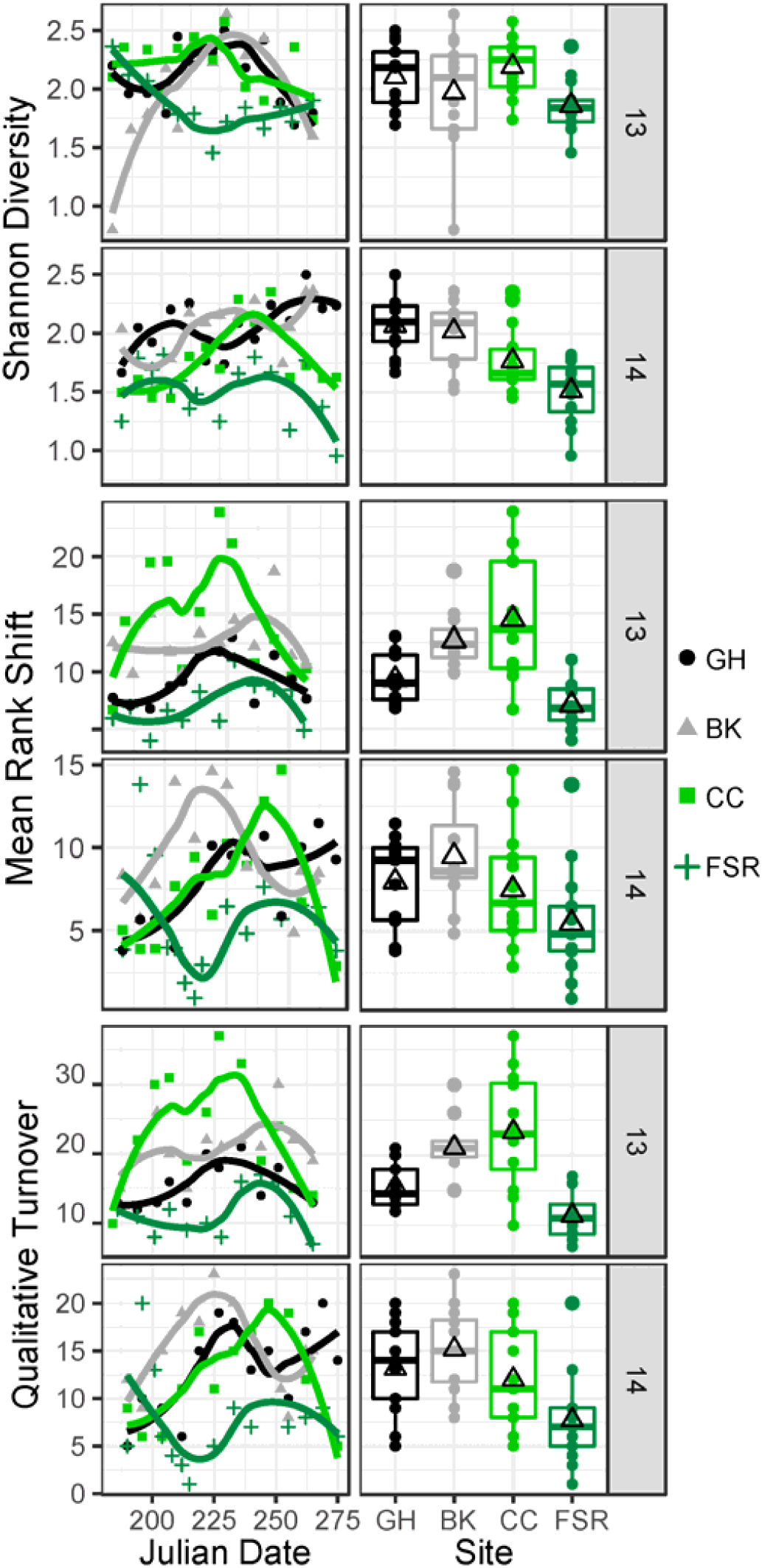
Time series and boxplots of Shannon diversity, mean rank shift and qualitative turnover (appearances plus disappearances between consecutive samplings) for 2013 and 2014. Open triangles in boxplots represent distribution averages. Boxplots are arranged from most urban (GH) to most agricultural (FSR).

NMDS analysis revealed similarity in forage composition within sites across years (Figure 6). Moreover, the arrangement of sites along NMDS axis one reflected their positions on the urbanization gradient, with highly urban sites GH and BK roughly coinciding, followed by the mixed urban and agricultural site CC, and finally the strictly agricultural site FSR.

**Figure 6:**
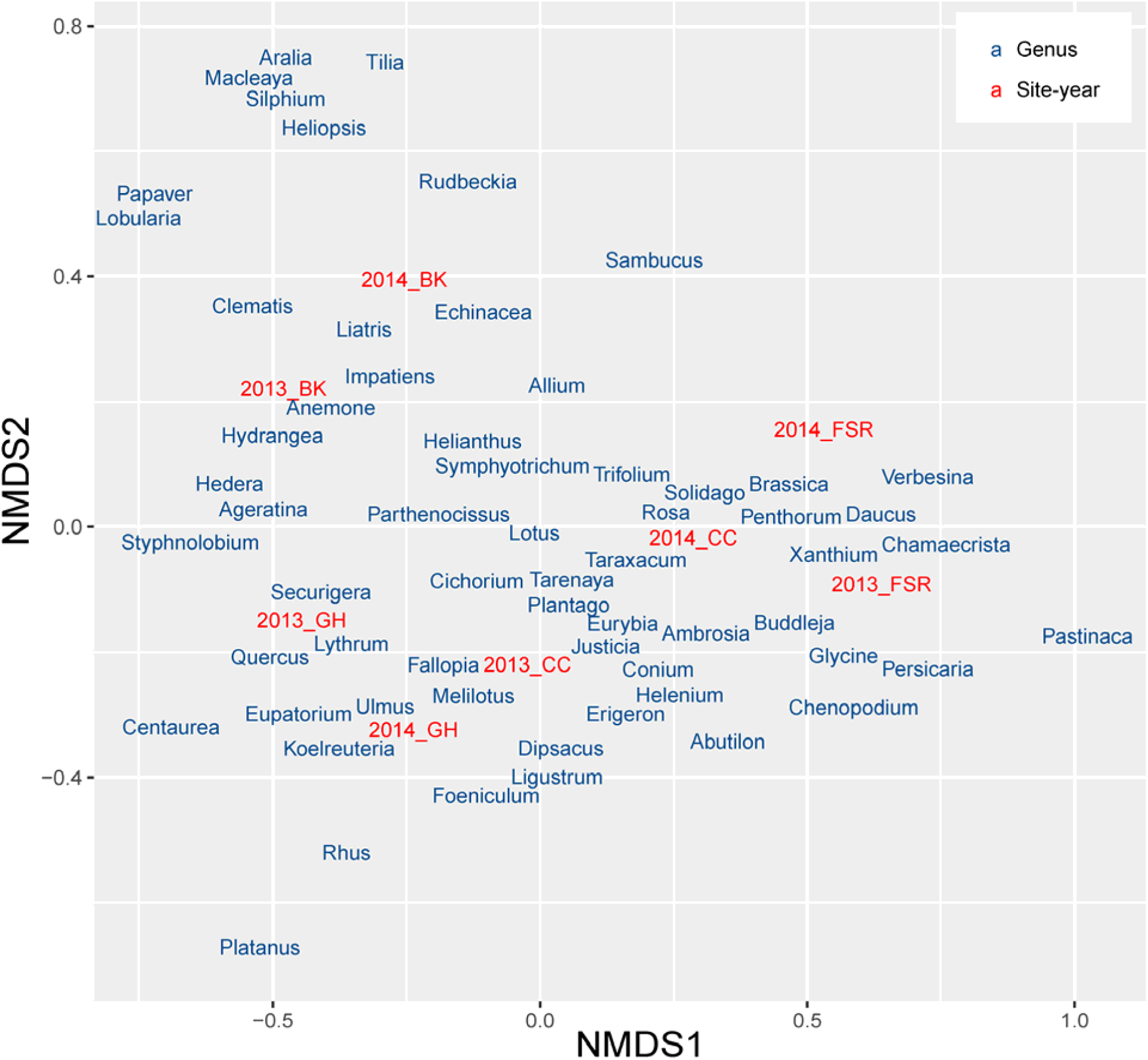
An NMDS biplot (k = 2, stress = 0.06) of forage compositional similarity between sites and years. Only genera with a maximum proportional abundance of at least 2.5 percent are plotted. Genus scores are plotted using weighted averages.

## Discussion

The most salient pattern in our data was that, consistent with our prediction, pollen diversity was greatest at the urban and suburban sites in our study system. It is important, however, not to jump to the interpretation that our more urban sites were more suitable for honey bees than the agricultural sites. To the contrary, prior studies in this system found that urban land use was negatively associated with colony productivity (Sponsler and Johnson 2015) and that honey bees situated at the agricultural-suburban divide of site CC dedicated significantly more foraging activity toward rural agricultural landcover components (Sponsler et al. 2017). While Ohio urban landcover apparently contains a diverse floral assemblage, a trait which likely makes it amenable to generalist solitary bees and bumble bees (Samuelson et al. 2018, Baldock et al. 2019), it may be depauperate in the high-volume floral resource patches which drive honey bee colony growth. Thus, future research contributions which seek to explicitly characterize the nutritional currency value of honey bee forage diversity may benefit from considering the implications of urban landscapes through the lens of optimal foraging theory and the life history of the honey bee. Investigations of whether elevated urban forage diversity is associated with decreased resource volume and patchiness is particularly warranted.

To the degree that honey bee foraging preferences are highly generalized, colonies containing thousands of foraging individuals enable efficient sampling of the surrounding floral community. The composition of the pollen samples collected from this study highlight the unique floral communities present in different landcover types. Particularly with respect to urban and suburban landscapes, small and heterogeneously managed parcels of land appear to result in greater overall plant community diversity relative to rural agricultural landscapes, at least for the spatial scales corresponding to honey bee foraging. This observation is consistent with the insight of Aronson et al. (2015), which suggests that the effect of urbanization on biological diversity depends strongly on the scale of investigation. At the scale of individual patches, urban land use may be associated with lower diversity due to the dominance of invasive species or the stringent filtering of species traits. At the opposite extreme of scale, geographically distant cities may exhibit low beta diversity due to the characteristic urban ecological conditions shared by all cities. Within cities, however, or along urban-rural gradients, urban land use may be associated with increased inter-patch biological diversity due to the extreme heterogeneity of current and historical land use found in urban areas. We suggest that the inter-patch scale of honey bee foraging combined with the inter-patch diversity of urban landscapes explains the increased diversity of pollen samples collected at our urban sites.

It is evident in Figure 4 that certain genera occur in our system as characteristically urban or agricultural taxa, while others are ubiquitous across the whole landscape gradient. The most striking example of the latter is *Trifolium*, which was found in high abundance in every pollen sample. *Plantago*, while less dominant, was similarly ubiquitous across sites. In both cases, this reflects the commonness of these genera both in agricultural interstices and in urban lawns and roadsides. In contrast, *Chamaecrista* was found almost exclusively in FSR and CC, sites partly or entirely agricultural. Conversely, *Hedera*, a naturalized ornamental that has become widespread in urban greenspaces, was found almost exclusively in the urban sites BK and GH. Similarly, *Hydrangea* is a frequently cultivated ornamental, and was found mainly at sites BK and GH. These patterns highlight the joint importance of incidental and intentional effects of human land use on floral community composition.

Humans have mediated fundamental transitions in the distributions and abundances of both managed and non-managed species (Achard et al. 2002, Keeley 2002, Foley et al. 2005, Klimaszewski-Patterson and Mensing 2016). Such community compositional change continues at a rapid pace (Seebens et al. 2015, Tarolli 2016, Seebens et al. 2017) and is often driven broadly by landscape-scale changes (Tarolli et al. 2014, Otto et al. 2016). Understanding how alterations in landscape structure and composition affect the composition and functioning of communities is a central goal in applied ecology. Nutritional limitation is likely a bottom-up factor involved in shaping biological communities, especially with respect to pollinators (Samuelson et al. 2018, Baldock et al. 2019, Wood et al. 2019). Thus, a greater understanding of pollinator landscape and nutritional ecology is central to improving pollinator health. In applying improved molecular pollen identification methods to honey bee collected pollen, we show that local landscape composition alters forage diversity, composition and temporal dynamics along a land use gradient. These findings provide direction for future integration of foraging ecology, landscape ecology and nutritional ecology on a broader range of taxa and environmental gradients.

## Acknowledgements

The authors thank D. Denlinger and K. Goodell for laboratory space, C Kurkul for sample processing and N Douridas for assistance at the Molly Caren Agricultural Center. This work was funded by a Project Apis m.—Costco Honey Bee Biology Fellowship, an OARDC SEEDS grant and support provided by state and federal funds appropriated to The Ohio State University, Ohio Agricultural Research and Development Center (OHO01277).

## Data Accessibility

All sequence data produced in this work can be downloaded from NCBI SRA, BioProject PRJNA574766. Descriptions of the bioinformatic analysis, R and python code, reference sequence databases, and supplemental excel files of the taxonomic inferences and the sequencing coverage per library can be found at https://github.com/RTRichar/UrbanRuralPollen.

## Author Contributions

Conceptualization, DBS, RTR, RMJ; Investigation, GC, TE, CHL, RTR; Formal Analysis, RTR, DBS; Resources, RMJ, RTR; Writing – Original Draft, RTR, DBS; Writing – Review & Editing, RTR, DBS, RMJ, CHL, GC, TE; Visualization, RTR, DBS; Supervision, RMJ, RTR, DBS, CHL; Project Administration, RMJ; Funding Acquisition, RMJ, RTR, DBS.

